# TEtrimmer: a novel tool to automate the manual curation of transposable elements

**DOI:** 10.1101/2024.06.27.600963

**Authors:** Jiangzhao Qian, Hang Xue, Shujun Ou, Jessica Storer, Lisa Fürtauer, Mary C. Wildermuth, Stefan Kusch, Ralph Panstruga

## Abstract

Transposable elements (TEs) are repetitive DNA sequences capable of moving within genomes. Accurate annotation and classification of TEs is crucial but challenging due to their sequence diversity and often fragmented occurrence. We present TEtrimmer, a novel tool to automate manual TE curation. TEtrimmer integrates multiple sequence alignment (MSA) clustering, MSA sequence extension, MSA cleaning, TE boundary definition, and TE classification, and provides report plots and a graphical user interface (GUI) application to inspect and improve results. Benchmarked on the genomes of six organisms from various kingdoms of life, TEtrimmer consistently improved the identification of intact TEs compared to established tools.

## Background

Transposable elements (TEs) are selfish repetitive DNA elements that can move within host genomes. TEs were first discovered by Barbara McClintock in maize in 1948 (McClintock, 1950). Since then, TEs have been identified in all studied eukaryotic species where they occupy a large proportion of many genomes, such as around 45% of the human genome (International Human Genome Sequencing Consortium et al., 2001), 53% of the zebrafish genome (Howe et al., 2013), and 85% of the maize genome (Schnable et al., 2009).

TEs were long regarded as “junk” DNA. However, TEs play key roles in genome evolution, development, and immunity (Bourque et al., 2018). For instance, TEs can regulate promoter and enhancer sequences to alter host gene expression (Fueyo et al., 2022). In addition, TEs are a source of many regulatory genes, including long non-coding RNAs (Kapusta et al., 2013) and small RNAs (Kunz et al., 2024). Moreover, TEs can be drivers of evolutionary innovations. A recent example of TE-driven evolution is an *Alu* element insertion into an intron of the human *TBXT* gene, which appears to have contributed to tail-loss evolution in primates (Xia et al., 2024).

TEs are classified according to their transposition mechanism. Class I encompasses the retrotransposons, which copy themselves for genome insertion via an RNA intermediate. Class I retrotransposons mainly include three subclasses, LTR (long terminal repeat) retrotransposons, LINEs (long interspersed nuclear elements), and SINEs (short interspersed nuclear elements) (Bourque et al., 2018; Wicker et al., 2007). An intact LTR retrotransposon always contains flanking terminal repeat sequences, while LINEs and SINEs are defined by their terminal genomic poly-A repeats. Class II, on the other hand, comprises DNA transposons, which do not usually generate copies but rather excise themselves for insertion into another locus in the host genome. The most prominent DNA transposons include TIR (terminal inverted repeat) and Helitron elements (Wicker et al., 2007). However, precise TE annotation remains challenging, mainly due to TE type divergence and the existence of decayed, fragmented, nested, and low-copy TEs, especially in genomes with a high TE content (Hoen et al., 2015; Storer et al., 2022).

TE characterization involves two main steps, which are TE discovery and TE annotation (Storer et al., 2022). Due to the repetitive nature of TEs, TE families can be represented by the TE consensus sequence library, also called TE library, which is constructed based on a multiple sequence alignment (MSA) of the TE copies from the respective reference genome (Goubert et al., 2022). The process of TE library construction is called TE discovery. The TE Hub Consortium (The TE Hub Consortium et al., 2021) has collected 51 tools to date (April 2024) that are associated with TE library construction (Sierra & Durbin, 2024). The term TE annotation refers to the process of genome-wide TE identification to correctly define both intact and fragmented TE copies in the genome. Pre-constructed TE libraries are available for some species from repositories such as RepBase (Weidong. Bao et al., 2015), Dfam (Hubley et al., 2016), and RepetDB (Amselem et al., 2019). To date, RepBase contains the most comprehensive collection of manually curated high-quality TE libraries, but these have been behind a paywall since 2018.

TE annotation is typically based on three main strategies, i.e., repository-based, repeat-based, and structure-based methods. The most-used strategy is the repository-based approach (Loreto et al., 2023), which takes advantage of public pre-constructed TE libraries for whole-genome TE annotation. Since closely related species share a common origin and exhibit genome sequence similarity, the established TE library can be used to identify homologous TEs in related genomes as well (Gao, 2023). This strategy heavily relies on the availability and quality of a suitable TE library for the species of interest. However, manually curated high-quality TE consensus libraries are only available for a limited number of species after years of community efforts (Hubley et al., 2016; Osmanski et al., 2023). In addition, because the current versions of RepBase are only available via paid subscription, many researchers, particularly from smaller institutes and/or less wealthy countries, do not have access to suitable TE libraries. By contrast, the repeat-based strategy generates a custom TE library from the genome of interest. This strategy mainly includes self-comparison and *k*-mer seeding methods (Storer et al., 2022). The self-comparison method identifies TEs by computationally intensive pairwise alignments between genomic sequences using algorithms such as RECON (Zhirong. Bao & Eddy, 2002) and Grouper (Quesneville et al., 2003). On the other hand, *k*-mer seeding methods like RED (Girgis, 2015), RepeatScout (Price et al., 2005), and P-Clouds (Gu et al., 2008) are generally faster and can efficiently handle large genomic datasets. However, the accurate clustering of TE sequences remains a challenge, and this method relies on high-quality genome assemblies. Poor genome quality can lead to inaccurate TE boundary definition, over-fragmentation of TE copies, and loss of TEs in incomplete assemblies (Kurtz et al., 2008). Finally, the structure-based strategy can annotate TEs by recognizing specific structural hallmarks such as LTRs, TIRs, target site duplications (TSDs), and terminal genomic poly-A repeats. A variety of algorithms, including LTR_FINDER (Xu & Wang, 2007), LTR_STRUC (McCarthy & McDonald, 2003), LTRharvest (Ellinghaus et al., 2008), TIR-Finder (Gambin et al., 2013), and SINE-Finder (Wenke et al., 2011) facilitate structure-based TE annotation. The structure-based methods can efficiently annotate low-copy TEs and precisely define TE boundaries; however, a high false-positive rate is associated with these approaches.

No single strategy or algorithm can comprehensively annotate TEs in a genome. Accordingly, various tools for de novo TE identification have been developed that integrate several annotation strategies, such as RepeatModeler2 (Flynn et al., 2020), EDTA2 (Ou et al., 2019) (https://github.com/oushujun/EDTA/tree/EDTA2), and REPET (Flutre et al., 2011). However, even these tools do not approach the gold standard quality of manually curated TE libraries (Platt et al., 2016).

Manual curation of TEs can be divided into five major steps: BLAST (Basic Local Alignment Search Tool), extend, extract, align, cluster, and trim (Goubert et al., 2022; Storer et al., 2021). In brief, the TE candidate from a de novo discovery pipeline is used for a nucleotide BLAST (BLASTN) query against the reference genome, and the coordinates of search hits are arbitrarily extended at both ends to include TE boundaries. Then, the TE sequences are extracted based on the extended coordinates and subsequently aligned. Finally, the sequences within the MSA are manually clustered based on their sequence relatedness and trimmed by removing regions of low conservation (Goubert et al., 2022; Platt et al., 2016; Storer et al., 2021). The manual curation of TEs can yield a gold standard custom TE library for the genome of interest but is very time-consuming to establish, and requires TE specialists with an in-depth understanding of TE structure and biology (Orozco-Arias et al., 2023). Accordingly, the comparability and reproducibility of TE libraries from manual curation may be limited by the level of expertise of the curators (Baril et al., 2024).

Some software tools, like EarlGrey (Baril et al., 2024) and MCHelper (Orozco-Arias et al., 2023), have been developed to partially automate and assist manual curation of TEs. EarlGrey mainly combines RepeatModeler and TEstrainer (https://github.com/jamesdgalbraith/TEstrainer) for TE library construction. TEstrainer is the core module to automate most manual curation steps but not the clustering of MSAs. TEstrainer attempts to bypass the MSA clustering by only including the top 20 BLASTN hits, which is insufficient for highly divergent TEs. MCHelper includes an MSA clustering method but lacks effective MSA cleaning ability. In essence, both EarlGrey and MCHelper can improve the precision of TE annotation in any given organism but do not yet approach the quality of manually curated TEs. Here, we introduce a novel user-friendly tool called TEtrimmer that automates the manual curation of TEs. TEtrimmer can efficiently cluster MSAs, clean MSAs, and precisely define TE boundaries. TEtrimmer provides comprehensive report plots and a graphical user interface (GUI) application to support the rapid inspection and improvement of results, which closes the gap to manual curation.

## Results

### Overview of TEtrimmer

Manual curation is crucial for a high quality of TE annotation. However, curation can involve hundreds to thousands of TE consensus sequences for each genome and is therefore time-consuming and requires expert TE knowledge. TEtrimmer is a software designed to automate the manual curation of TEs. The input of TEtrimmer can be the output TE consensus libraries from de novo TE discovery tools like EDTA2, RepeatModeler2, and REPET and/or TE libraries from closely related species. To allow multi-thread analysis, TEtrimmer separates each set of input sequences into a single file (Fig. 1). For each separated sequence, TEtrimmer automatically performs (BLASTN searches against the respective genome (Ye et al., 2006). It uses the retrieved BLAST hits for MSA generation, MSA clustering (Fig. 2), MSA sequence extension, and MSA cleaning (Fig. 3).

**Figure 1:**
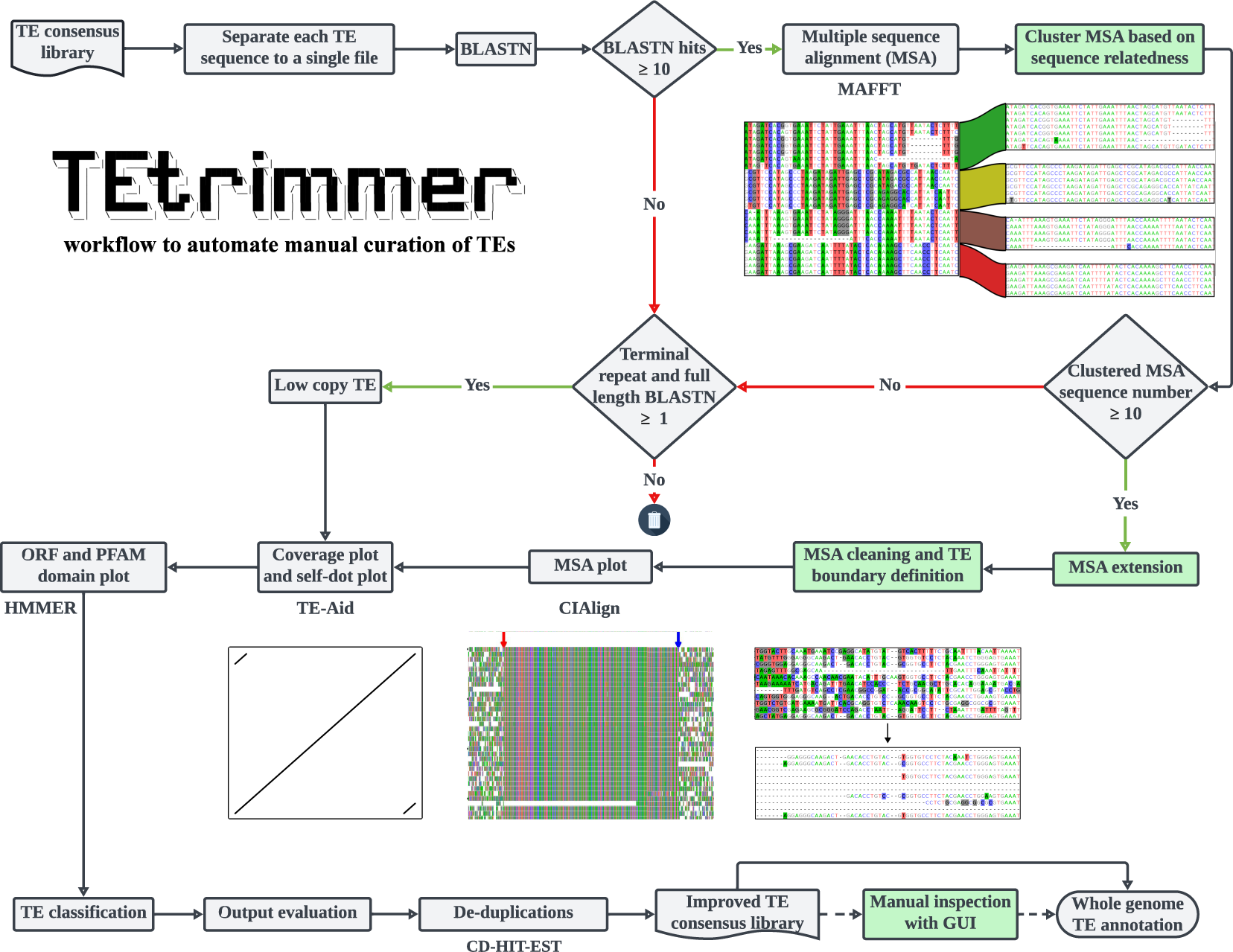
TEtrimmer workflow. TEtrimmer is a multi-thread tool to automate the manual curation of TEs. It can perform BLASTN searches of TE sequences retrieved from any TE consensus library against the respective genome and facilitate the generation of MSAs, MSA clustering, MSA sequence extension, MSA cleaning, TE boundary definition, and TE classification. Moreover, TEtrimmer provides detailed report plots and offers a graphical user interface (GUI) application to enable inspection and improvement of each output. The key features of TEtrimmer are highlighted by the green background color.

**Figure 2:**
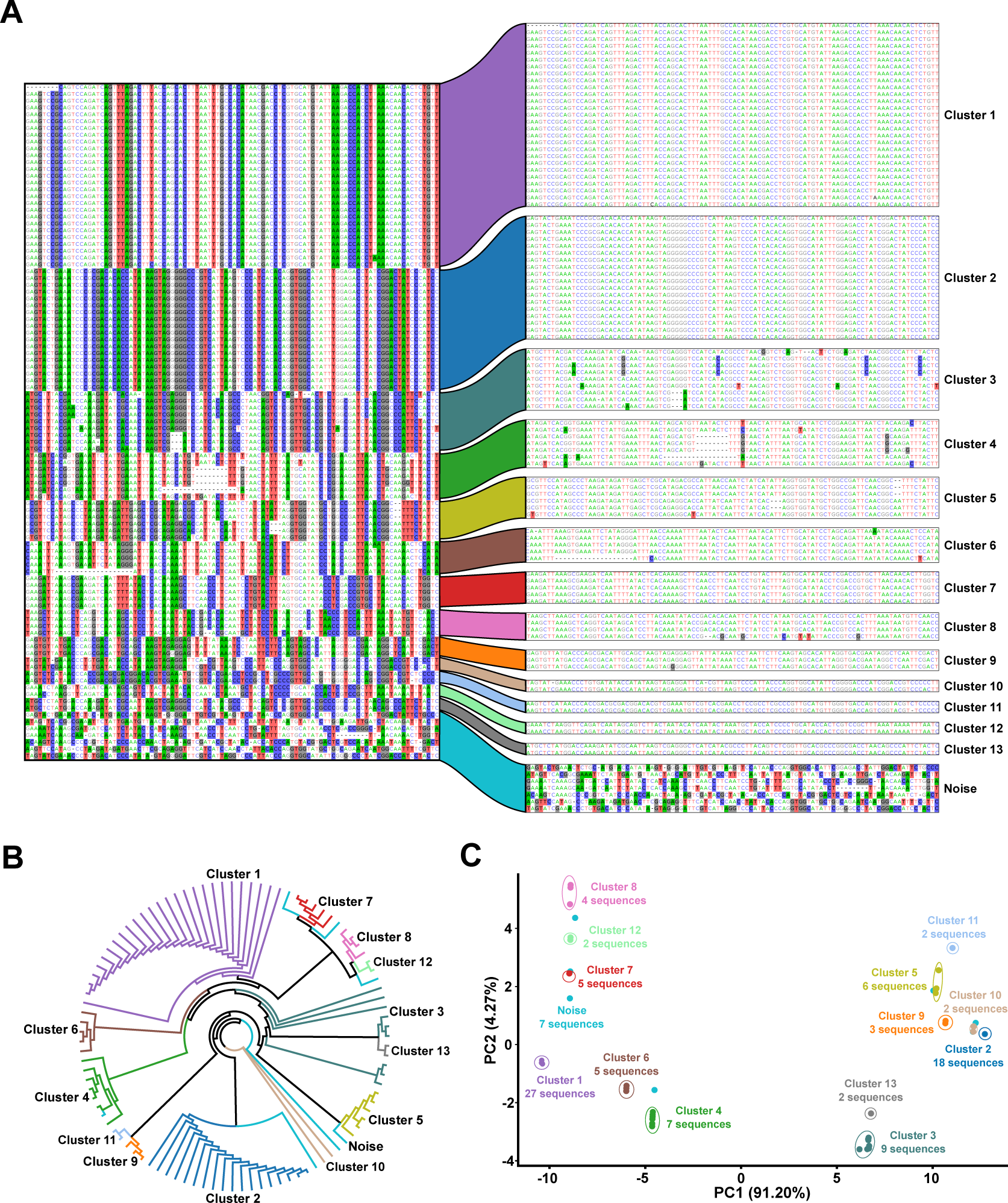
TEtrimmer efficiently clusters sequences within the MSA. One *D. melanogaster* LTR retrotransposon (named rnd-1-family-39), identified by RepeatModeler2, was chosen as an example for further analysis. TEtrimmer performed a BLASTN search of this LTR retrotransposon against the *D. melanogaster* genome, established an MSA with the respective BLASTN hit sequences, and conducted MSA clustering. **A** The left part represents the divergent columns in the MSA before clustering. The right area displays the MSAs after clustering by TEtrimmer based on the phylogenetic tree (B) relative branch distance-based DBSCAN analysis (epsilon = 0.1, min sample number = 2). Nucleotide background colors in the MSA represent sites where the proportion of the respective nucleotide in that column is below 0.4. The cluster names given here were consistently used in (B) and (C). Each color represents the corresponding cluster. **B** The un-rooted maximum likelihood phylogenetic tree generated by IQ-TREE is based on the MSA before clustering. Cluster labels and colors are consistent with (A). **C** PCA plot calculated using the phylogenetic tree relative branch distance matrix (B). The percentage of variance explained by the respective principal components PC1 and PC2 is indicated on the x-and y-axis, respectively. Cluster labels and colors correspond with (A). The number of sequences within each cluster is indicated.

**Figure 3:**
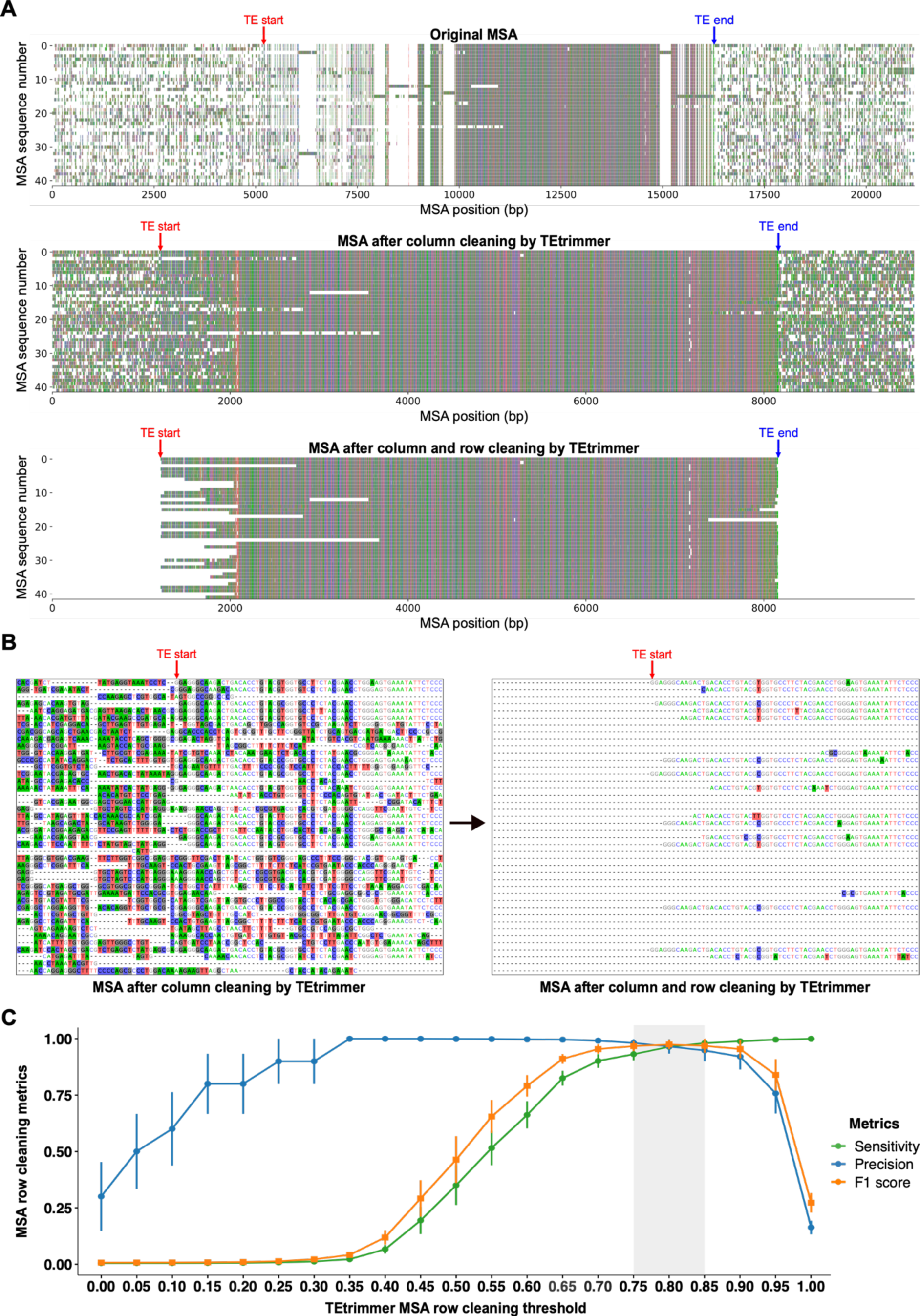
TEtrimmer succeeds in removing lowly conserved regions from MSAs. A *B. hordei* LINE element (named rnd-1-family-34), identified by RepeatModeler2, was chosen as a representative example for demonstrating the TEtrimmer MSA cleaning performance. **A** After BLASTN search of the selected LINE sequence against the *B. hordei* genome, sequence extension, and MSA generation, the LINE boundaries were identified, which are indicated by “TE start” (red) and “TE end” (blue). Nucleotides are represented with colored bars; gaps are indicated as blank regions in the plots. The top panel is the original MSA before cleaning, which contains many gappy columns and noisy rows. After MSA column cleaning by the TEtrimmer function “`remove_gap_columns`”, the majority of the gappy columns are removed (middle panel). Then, TEtrimmer cleans sequences in the MSA row by row using the TEtrimmer function “`crop_end_by_divergence`” (bottom panel), which removes lowly conserved regions (bottom panel). **B** The left and right panels are both the magnified MSA regions near the selected LINE element left boundary. The boundary is indicated by “TE start” (red). The left panel illustrates the MSA before and the right panel after row cleaning by TEtrimmer via “`crop_end_by_divergence`”. Nucleotide background colors in the MSA represent sites where the proportion of the respective nucleotide is below 0.4. **C** Ten TEs were randomly selected from the RepeatModeler2 consensus libraries of *B. hordei* (barley powdery mildew fungus), *D. melanogaster* (fruit fly), *D. rerio* (zebrafish), and *O. sativa* (rice). After BLASTN searches of the selected sequence against the corresponding genomes, sequence extension, and MSA column cleaning, the generated MSAs were used for benchmarking. Manual cleaning was performed to serve as a reference to enable assessment of TEtrimmer MSA cleaning performance. The MSAs were cleaned by the TEtrimmer cleaning function “`crop_end_by_divergence`” using cleaning thresholds ranging from 0 to 1 and a sliding window size of 40 bp (a detailed description for thresholds and sliding windows can be found in the Methods section). A confusion matrix analysis was conducted to evaluate the TEtrimmer cleaning performance. The x-axis shows the cleaning threshold used by the TEtrimmer function “`crop_end_by_divergence`”; the y-axis displays the confusion matrix score for the metrics sensitivity (green), precision (blue), and F1 score (orange, calculated by (2 * (sensitivity * precision) / (sensitivity + precision))). Standard error bars are shown in the plot. The grey-shaded box indicates the threshold range where all three metrics scores are above 0.93.

Thereafter, TEtrimmer employs the clustered, extended and cleaned MSAs to generate consensus sequences for the definition of putative TE boundaries. Then, potential terminal repeats are identified, and a prediction of open reading frames (ORFs) and protein domains on the basis of the protein families database (PFAM) (Mistry et al., 2021) are conducted. Subsequently, TE sequences are classified and an output evaluation is performed mainly based on the existence of terminal repeats, and the full length BLASTN hit numbers (Table 1). Consequently, two TE consensus library files are produced by TEtrimmer. The first file contains all output sequences while the second file is generated by removing duplicated sequences via CD-HIT-EST (Fu et al., 2012) (Fig. 1; see Methods for further details).

**Table 1.** Criteria for evaluation levels of the TEtrimmer output ^a^.

| Category | Presence of terminal repeats | Classified | Sequence number in the MSA | Number of full-length BLASTN hits | Presence of PFAM domain |
| --- | --- | --- | --- | --- | --- |
| Perfect | TRUE | TRUE | ≥30 | ≥5 | TRUE |
| Good | TRUE | Not_required | ≥10 | ≥2 | Not_required |
| Reco_check | Not_required | Not_required | ≥20 | ≥2 | Not_required |
| Need_check | Not_required | Not_required | ≥10 | Not_required | Not_required |
| Low_copy | TRUE | Not_required | < 10 | ≥2 | Not_required |

<sup>a</sup> These criteria might not efficiently evaluate TEs lacking terminal repeats such as LINEs and SINEs.

TEtrimmer provides a series of report plots for each output sequence to enable detailed inspection (Fig. 4). Users can choose to use the TEtrimmer-generated TE consensus library directly for genome-wide TE annotation or take advantage of the supplied GUI application (Fig. 5) to inspect and curate the TE library, which can generate a gold standard manual curation-level TE library. A detailed description of each step can be found in the Methods section.

**Figure 4:**
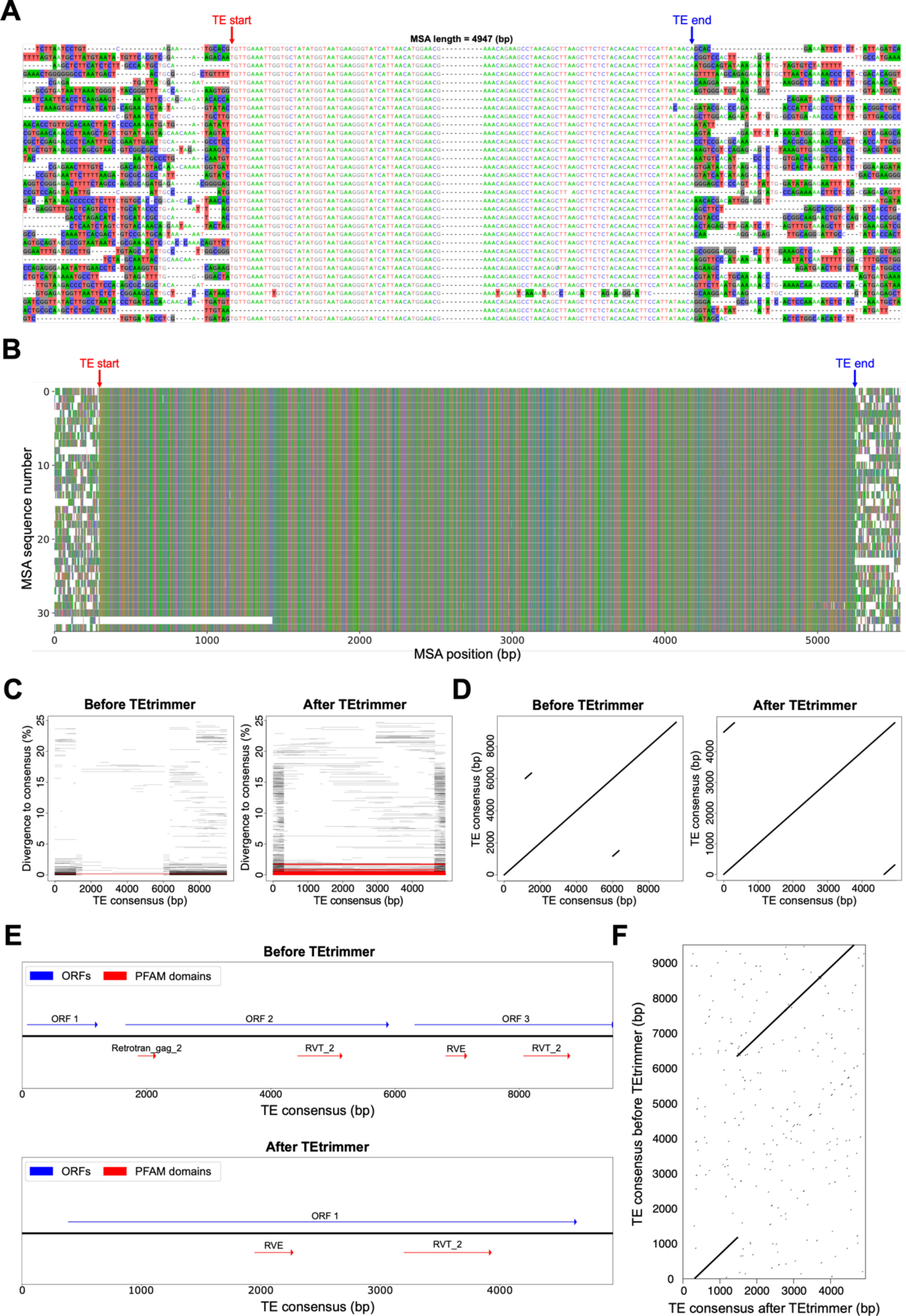
TEtrimmer provides comprehensive report plots for each discovered TE. The *B. hordei* LTR retrotransposon named “ltr-1-family-22”, initially annotated by RepeatModeler2 and re-analyzed with TEtrimmer, is shown as an example. **A** The alignment of 33 sequences in total shows the TE boundary regions of the MSA (100 nucleotide sites are displayed for each side) after TEtrimmer analysis. Nucleotide background colors in the MSA represent sites where the proportion of the respective nucleotide is below 0.4. The TE boundaries defined by TEtrimmer are indicated as “TE start” (red) and “TE end” (blue). The two boundary regions were artificially connected by ten gaps (“-”) and the total length of the MSA is indicated on the top. **B** The plot shows the entire MSA after TEtrimmer analysis. The TE boundaries are indicated as in (A). Nucleotides are represented with colored bars; gaps are indicated as blank regions in the plot. The y-axis gives the nucleotide position (in bp) within the MSA. **C** The panels show a BLASTN plot before (left panel) and after (right panel) TEtrimmer analysis, each following a BLASTN search of the respective TE consensus sequence to identify TE sequence hits at a genome-wide scale. The x-axis indicates the TE consensus nucleotide position (in bp) and the y-axis is the sequence divergence in percent compared to the TE consensus sequence. Each line indicates a BLASTN hit; red lines highlight hits with a sequence divergence below 1.5% and a sequence coverage > 90%. **D** Self-dot plots before (left panel) and after (right panel) TEtrimmer analysis. The axes show the nucleotide position (in bp) in the TE consensus sequence. The repeat regions are represented by short diagonal lines outside the main diagonal. **E** The plot shows a genomic map of open reading frames (ORFs; blue arrows) and PFAM domains (red arrows) predicted in the TE consensus sequence before (upper panel) and after (lower panel) TEtrimmer analysis. The direction of the arrows indicates the strands of the respective ORFs/PFAMs in the sequence. ORF and PFAM identity are indicated on top of the arrows. After TE trimmer analysis, ORF2 is removed and ORF1 and ORF3 are fused to make the new ORF1. **F** Dot plots (window size = 25 bp, threshold = 50; the threshold represents the minimal sum of substitution scores in a defined window required for a dot to be plotted) of the TE consensus sequence before (y-axis) and after (x-axis) TEtrimmer analysis. Axes show the nucleotide position (in bp). Identical regions are represented by short diagonal lines outside the main diagonal.

**Figure 5:**
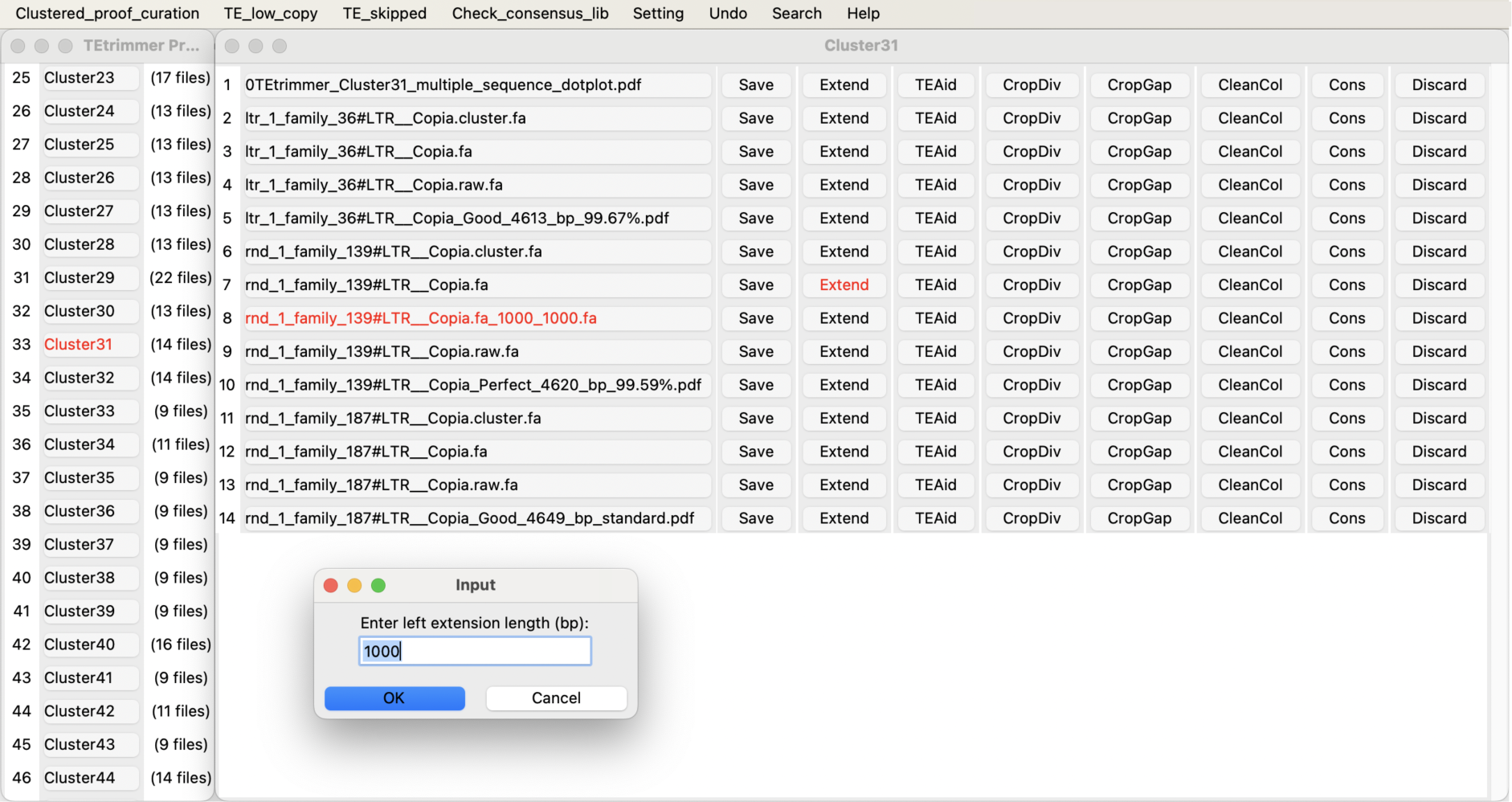
TEtrimmer provides a user-friendly graphical user interface (GUI) application to inspect and improve discovered TEs. Four files are associated with each consensus sequence (see main text for further details). All files are grouped together when their corresponding consensus sequences have >90% identity (“Cluster”). Users can inspect each result by selecting the corresponding file name. TE sequences in the MSA can be extended by activating the “Extend” button. The “TEAid” button helps to generate the interactive report plot (see Fig. 4). The buttons “CropDiv”, “CropGap”, and “CleanCol” relate to the MSA cleaning functions “`crop_end_by_divergence`”, “`crop_end_by_gap`”, and “`remove_gap_columns`”, respectively. The button “Cons” can be used to generate a consensus sequence based on the corresponding MSA.

### TEtrimmer efficiently clusters sequences in the MSA

The TEtrimmer tool initially performs a BLASTN search for each input TE consensus sequence against the corresponding genome (Fig. 1). If more than ten hits are obtained, an MSA is established; otherwise the sequence is considered a potential low-copy TE. Due to the often high sequence similarity among TE sub-families, this initial BLASTN search typically retrieves several types of TE sub-families per input sequence. These can be visualized by distinct patterns of the aligned TE sequences corresponding to the BLASTN hits (Fig. 2A). Thus, the first labor-intensive task of the traditional manual curation of TEs is to separate TE sub-families within the MSA according to sequence relatedness. TEtrimmer combines a phylogenetic tree approach (Fig. 2B) and a machine learning method (Density-Based Spatial Clustering of Applications with Noise, DBSCAN) (Birant & Kut, 2007) to separate automatically different alignment patterns that might correspond to distinct TE sub-families.

To illustrate the TEtrimmer clustering ability, one LTR retrotransposon from *Drosophila melanogaster* identified by RepeatModeler2 was selected as an example. The BLASTN search of the selected TE sequence against the *D. Melanogaster* genome yielded 99 hits. After extraction of the BLASTN hit sequences and generation of an MSA, 101 divergent columns were selected in the original MSA and combined into a new MSA of pseudo-sequences representing the divergent regions in the original MSA (Fig. 2A, left part). Divergent columns are defined as columns where the proportion of the predominant nucleotide in that column is below 0.8. The combined MSA based on the divergent columns was used to construct an un-rooted maximum-likelihood phylogenetic tree (Fig. 2B), which was employed to calculate a relative tree branch distance matrix (Supplementary Table 1). Finally, DBSCAN used the distance matrix for MSA clustering, which yielded fourteen sub-MSAs (“clusters”), likely corresponding to different TE subfamilies in the original MSA (Fig. 2A, right part and Fig. 2B). Thirteen of these clusters, cluster 1 to 13, had clearly grouped sequences. The fourteenth cluster was dubbed “noise”, which means the sequences in this cluster could not be sub-grouped further. The TEtrimmer MSA clustering efficiency was additionally supported by a principal component analysis (PCA) plot based on the phylogenetic tree relative branch distance matrix (Fig. 2C). In the case of clusters 1 to 13, the sequences in each cluster grouped adjacent to each other but were separated from the sequences of other clusters. Regarding the “noise” cluster, its seven sequences were randomly distributed across the entire PCA plot, indicating their low relatedness. This rather intricate example of a complex MSA pattern illustrates that TEtrimmer can efficiently cluster TE sequences in an MSA.

### TEtrimmer efficiently extends and precisely cleans clustered MSA sequences

As elements retrieved by BLASTN searches can be incomplete, the sequences in the clustered MSAs typically need to be extended. TEtrimmer offers a convenient function for the expansion of sequence ends in the MSAs, which is iteratively used by the tool until the sequences cover the putative TE boundaries (Fig. 1; see Methods for further details on the sequence extension procedure). However, sequence extension beyond the actual TE borders usually results in non-TE sequences added to the MSA. These non-TE sequences, being typically very divergent, consistently produce badly aligned “noisy” regions in the MSA (Fig. 3A). Cleaning these regions is a time-consuming step of the manual curation process. The cleaning (“trimming”) module of TEtrimmer efficiently addresses this task by removing gappy columns and the majority of lowly conserved regions within rows of MSAs after sequence extension. We demonstrate this capability by a representative MSA based on extended sequences derived from a *B. hordei* LINE element, rnd-1-family-34, (Fig. 3A). After MSA column cleaning by the TEtrimmer function “`remove_gap_columns`”, the majority of gappy columns within the original MSA were eliminated (top and middle panel of Fig. 3A). Subsequently, the remaining noisy regions of the MSA were cleaned row by row by the TEtrimmer function “`crop_end_by_divergence`” (middle and bottom panel of Fig. 3A and B). Following column and row cleaning of the MSA, TE boundaries (“TE start” and “TE end”) are determined by TEtrimmer (Fig. 3A and B, see Methods for details).

To evaluate the MSA cleaning performance, we compared manual cleaning to TEtrimmer-based cleaning for the above-mentioned LINE element. The effectiveness of cleaning conducted by TEtrimmer was assessed quantitatively using a customized confusion matrix (Provost et al., 1998), which showed a cleaning sensitivity of 0.977 and a precision of 0.999. This means that TEtrimmer correctly identified and removed 97.7% of the manually cleaned MSA regions and 99.9% of the regions it removed were in agreement with manual cleaning. An additional MSA row cleaning function of TEtrimmer not illustrated here is “`crop_end_by_gap`”. In the TEtrimmer analysis process, this is an additional option preferentially used for LINE elements to deal in particular with their highly divergent 5’ regions. In contrast to the “`crop_end_by_divergence`” function, which is based on nucleotide proportions, “`crop_end_by_gap`” takes advantage of gap information for the cleaning process (see Methods for further details).

To test the cleaning performance across various types of TEs and different species, we randomly selected ten TEs from *B. hordei*, *D. melanogaster*, *D. rerio*, and *O. sativa*, including LTR retrotransposons, LINEs, and DNA transposons. After conducting BLASTN searches of the selected TE sequences against the corresponding genomes, sequence extension, MSA generation, and removal of gappy columns, we performed both manual and TEtrimmer-based MSA cleaning, the latter using the function “`crop_end_by_divergence`” with different thresholds. A detailed explanation of the cleaning thresholds can be found in the Methods section. TEtrimmer showed optimal row cleaning performance with a set threshold between 0.75 and 0.85, yielding averaged sensitivity and precision scores above 0.959 and 0.971, respectively (Fig. 3C). The results highlight the excellent MSA cleaning ability of TEtrimmer across various TE types from different species. Accordingly, the software name “TEtrimmer” relates to the advanced MSA trimming ability of the tool.

### TEtrimmer provides detailed report plots and summary tables for each discovered TE

Detailed report plots and summary tables for each output facilitate the assessment of outputs and highlight differences in TE consensus sequences before and after TEtrimmer analysis (Fig. 4 and Supplementary Table 2). A *B. hordei* LTR retrotransposon, named “ltr-1-family-22” and initially identified by RepeatModeler2, was used for illustration of the report plots (Fig. 4). The first plot presents the MSA columns around TEtrimmer-defined TE boundaries. The precise boundary positions are denoted by colored arrows (Fig. 4A). This plot can help to inspect the existence of TSDs and the accuracy of TE boundary definition. In the example, the marked boundaries clearly separate the highly divergent from the conserved regions and TSD sequences can be identified (e.g. “GCACG” for the first sequence in Fig. 4A), which represents a correct boundary definition as judged by manual inspection. While the TE boundary plot only shows the ends of the MSA, the second plot displays the entire MSA (Fig. 4B), which enables users to verify the alignment quality of the full MSA after TEtrimmer analysis. The third type of plot illustrates the divergence of BLASTN hits against the TE consensus sequence and their coordinates along the TE consensus sequence (Fig. 4C). For instance, the selected exemplary LTR retrotransposon initially showed only one full-length BLASTN hit (Fig. 4C left panel), but 77 full-length BLASTN hits were recovered after TEtrimmer analysis, which indicates a substantial improvement (Fig. 4C right panel). The fourth type of plot is a self-dot plot, which can reveal TE sequence features such as flanking LTRs and TIRs (Fig. 4D). The example self-dot plot after TEtrimmer analysis shows flanking LTRs from bp 1 to 310 and bp 4638 to 4947 of the output consensus sequence (Fig. 4D right panel), but only internal repeat regions (from bp 1167 to 1506 and bp 6003 to 6342), indicative of a faulty TE, were identified for the input consensus sequence (Fig. 4D left panel). Besides BLASTN hits and sequence features, ORF and protein domains are also crucial for evaluating a TE consensus sequence. TEtrimmer provides a fifth type of plot to display the ORF and PFAM domains of the consensus sequence before and after TEtrimmer analysis (Fig. 4E). For instance, after TEtrimmer analysis ORF 2 was eliminated by deleting the sequence region from bp 1167 to 6002 of the input consensus sequence. Additionally, ORF 1 and ORF 3 of the input consensus sequence (Fig. 4E, upper panel) were fused into the single ORF 1 of the output consensus sequence (Fig. 4E, lower panel). The final plot is a dot plot comparing TE consensus sequence similarity before and after TEtrimmer analysis (Fig. 4F). The dot plot highlights both similar and different regions between the two sequences. For example, the regions from bp 0 to 1166 and bp 6343 to 9508 of the input sequence were identical with bp 311 to 1477 and bp 1478 to 4637, respectively, of the output sequence. In addition to the mentioned plots, TEtrimmer provides a summary table compiling the analysis results such as the numbers of BLASTN hits, sequence lengths, TE classifications, and the existence of terminal repeats (Supplementary Table 2). The summary table helps to browse and index analysis results conveniently. By combining the report plots and the summary table, users can quickly evaluate the accuracy of TEtrimmer outputs and their differences from the input TE consensus sequence.

### TEtrimmer provides a user-friendly graphical user interface application for inspecting and improving discovered TEs

To examine and improve the de novo-generated TE consensus libraries established by TEtrimmer further, a GUI application is provided, which allows inspecting and correcting outputs conveniently. Four files are associated to each consensus sequence in the GUI: (1) the original MSA based on the BLASTN search of the input TE consensus sequence against the corresponding genome, (2) the MSA after clustering, sequence extension, and MSA cleaning but before TE boundary definition, (3) the MSA with defined TE boundaries used to generate the final TE consensus sequence, and (4) several report plots (Fig. 4). To group highly identical sequences, TEtrimmer calculates the nucleotide identity among all TE consensus output sequences. All four files mentioned above that relate to highly similar TE consensus sequences (identity >90%) are placed together into one “TE cluster” and listed by the GUI application (left window in Fig. 5).

For instance, after TEtrimmer analysis, the TE sequences “ltr_1_family_36”, “rnd_1_family_139”, and “rnd_1_family_187”, which share more than 99.59% identity, and their associated files were placed into one “TE cluster” (Cluster 31; right window in Fig. 5). Moreover, multiple sequence dot plot files were provided to help visualizing consensus sequence similarities within a “TE cluster” (Supplementary Fig. 1). The multiple sequence dot plots help users conveniently inspect and compare similar outputs. Clustered files can be accessed by selecting the corresponding file names. Users can quickly assess each consensus sequence via the corresponding report plots file (Fig. 4) (e.g. “rnd_1_family_139#LTR Copia_Perfect_4620_bp_99.59%.pdf”). If the TEtrimmer output appears to be inaccurate, users can easily modify and improve it using the provided MSA files and functions. For example, if the sequence extension is insufficient in the view of the user, it can be manually elongated by activating the “Extend” button (right window in Fig. 5). The default extension size is 1000 bp for both ends, and these values can be easily modified separately for each side. After extension, a new file will appear with the extended sequences (e.g. “rnd_1_family_139#LTR Copia.fa_1000_1000.fa”, with 1000 bp extension on each side). Furthermore, to assist in identifying TE boundaries after manual extension, we integrated plot functions into the GUI application, allowing users to generate interactive report plots (Supplementary File 1) by activating the “TEAid” button. Additionally, the buttons “CropDiv”, “CropGap”, and “CleanCol” can be used to clean the MSA efficiently, which represent the same functions used for automated MSA cleaning in the TE trimmer pipeline (Fig. 3). In conclusion, the TEtrimmer GUI application integrates various files and functions associated with sequence extension, MSA cleaning, boundary identification, and visualization of TE consensus sequences, enabling users to examine and improve the TE consensus library conveniently. With the TEtrimmer GUI, the manual curation level of the TE consensus library can be readily achieved.

### TEtrimmer improves the recovery of intact TEs compared to EDTA2 and RepeatModeler2

We tested the TEtrimmer performance on six organisms belonging to various kingdoms of eukaryotic life. These were *Blumeria hordei* (barley powdery mildew fungus), *Drosophila melanogaster* (fruit fly), *Danio rerio* (zebrafish), *Oryza sativa* (rice), *Zea mays* (maize), and *Homo sapiens* (human) (Table 2). Benchmarking was initially conducted by directly comparing reference TE consensus libraries with de novo-generated libraries derived from EDTA2, RepeatModeler2, and both these tools after additional TEtrimmer analysis (consensus library after de-duplication). For this approach, we adopted the benchmarking method reported by Flynn and co-workers (Flynn et al., 2020). Four benchmarking levels were assigned to evaluate sequence matches between the reference libraries and the de novo-generated libraries, including “Perfect”, “Good”, “Present”, and “Not found”. For the analysis, LTR retrotransposons were separated into flanking LTRs and LTR internal sequences (LTR-INT) for all de novo-generated and reference libraries. We found that up to 74% of the “Perfect” families were comprised of flanking LTRs (Supplementary Table 3). To mitigate the potential bias caused by these sequences, we eliminated all flanking LTRs for this benchmarking (see Methods for details).

**Table 2.**
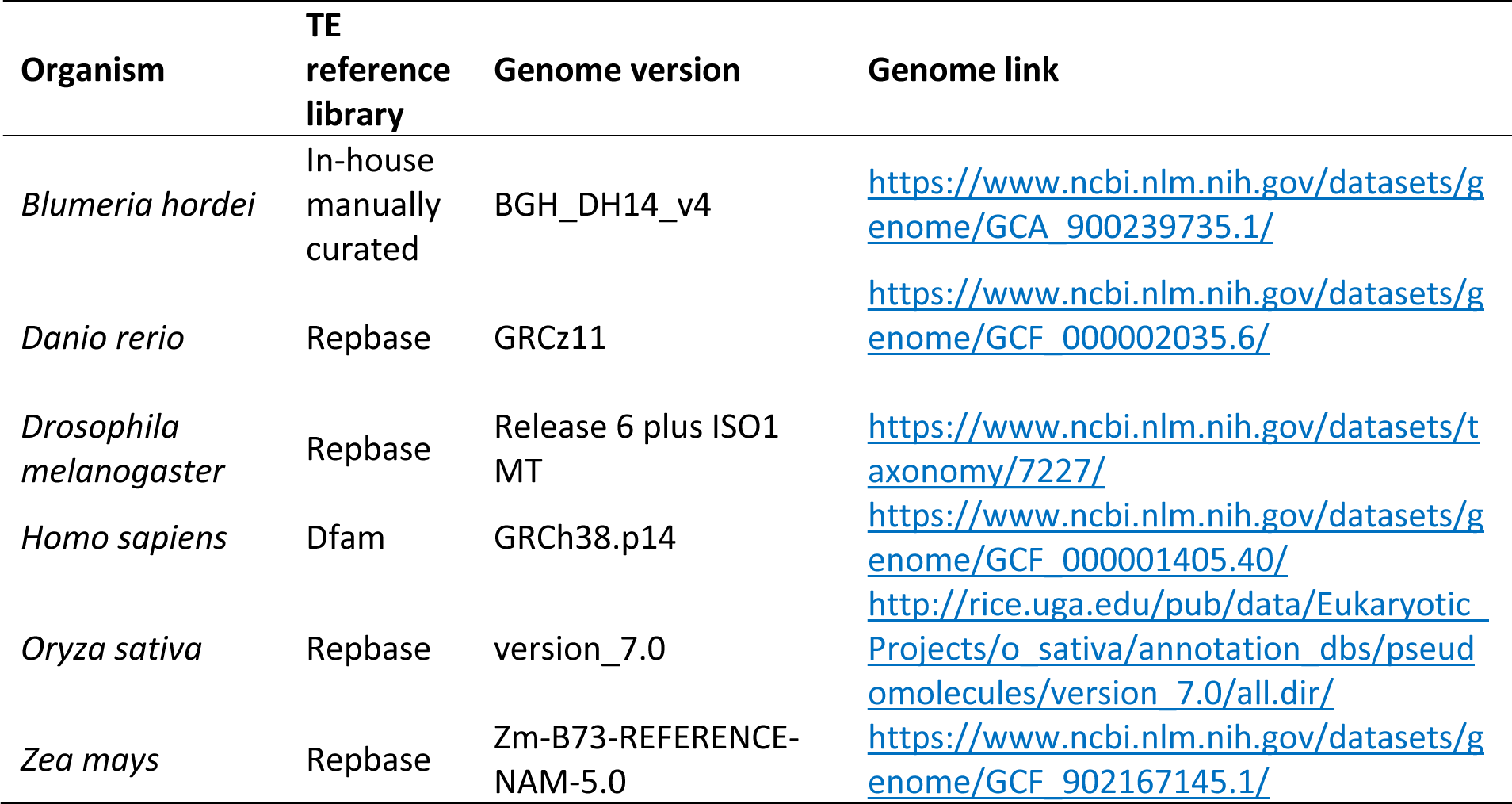
Genomes and TE reference libraries used to benchmark TEtrimmer.

The comparison between TEtrimmer results based on EDTA2 (EDTA2+TEtrimmer) and the EDTA2 consensus library (EDTA2 lib) revealed that TEtrimmer considerably improved the identification of “Perfect” transposable element (TE) families by producing 1.3-to 29.8-times more “Perfect” TE families across all six tested organisms (Fig. 6A). Specifically, EDTA2+TEtrimmer consistently generated more DNA and LINE elements in the “Perfect” family category but fewer LTR retrotransposons in case of *O. sativa* (Fig. 6B). EDTA2 struct+lib was generated by combining the EDTA2 consensus library with all TE sequences discovered by the EDTA2 structure-based module, followed by the removal of nearly identical sequences. EDTA2+TEtrimmer revealed more “Perfect” families than EDTA2 struct+lib for most of the selected species, except for *O. sativa* and *Z. mays*. TEtrimmer results based on RepeatModeler2 (RM2+TEtrimmer in Fig. 6A) generated 1.2-to 2.2-times more “Perfect” families and consistently identified more DNA and LINE elements as “Perfect” families across all selected organisms compared to RepeatModeler2 alone (RM2 in Fig. 6A). Conversely, fewer LTR retrotransposons were identified by TEtrimmer for *D. rerio*, *O. sativa*, and *Z. mays* (Fig. 6B and Supplementary Table 4), which might be due to the relatively inefficient identification of low-copy TEs by TEtrimmer. Overall, TEtrimmer predominantly improved the curation of TEs in all organisms except for LTR retrotransposons in the two plant genomes.

**Figure 6.**
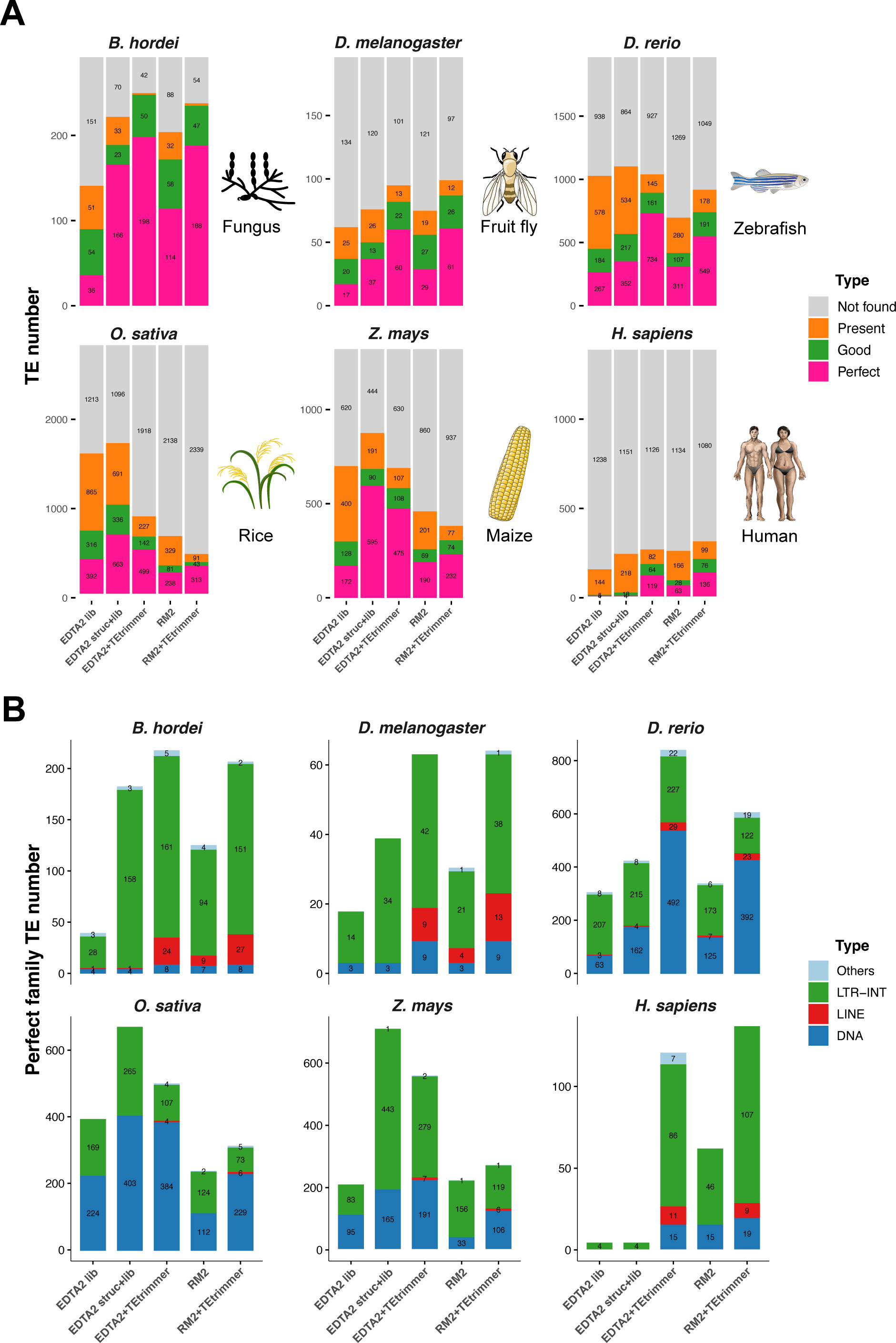
TEtrimmer improves the discovery of intact TEs compared to EDTA2 and RepeatModeler2. We benchmarked the performance of EDTA2, RepeatModeler2 (RM2), and both these tools after additional TEtrimmer analysis (EDTA2+TEtrimmer and RM2+TEtrimmer, respectively) by comparing the TE quality in the de novo-generated consensus libraries with the quality in the corresponding reference TE consensus libraries (Table 2) of the genomes of six organisms, i.e., *B. hordei* (barley powdery mildew fungus), *D. melanogaster* (fruit fly), *D. rerio* (zebrafish), *O. sativa* (rice), *Z. mays* (maize), and *H. sapiens* (human). Two libraries were used for EDTA2, termed “EDTA2 lib” and “EDTA2 struct+lib”. “EDTA2 lib” represents the EDTA2 consensus library and “EDTA2 struct+lib” was generated by combining the EDTA2 consensus library with all TE sequences discovered by the EDTA2 structure-based module. **A** The stacked bar graphs show the proportion of correctly discovered consensus TEs by the respective analysis tools/pipelines indicated on the x-axis according to the color code shown on the right. The y-axis displays the number of TE sequences in the respective reference TE library, i.e., 292 for *B. hordei*, 196 for *D. melanogaster*, 1967 for *D. rerio*, 2786 for *O. sativa*, 1320 for *Z. mays*, and 1391 for *H. sapiens*. Benchmarking categories included “Perfect” (pink), “Good” (green), “Present” (orange), and “Not found” (grey) (Flynn et al., 2020). “Perfect” means that the reference TE consensus sequences have one match in the de novo-generated library with > 95% similarity and coverage. “Good” indicates that the reference TE consensus sequence has multiple overlapping matches in the de novo-generated library, each with > 95% similarity and full coverage. “Present” is similar to “Good”, but the required minimum similarity and total coverage is decreased to 80%. The remaining TE sequences were assigned to the “Not found” category. **B** The stacked bar graphs show the number of “Perfect” matches in (A) resolved for the TE types of DNA transposons, LINEs LTR retrotransposon internal sequences (LTR-INT), and others (including e.g. Helitrons, SINEs, and simple repeat elements). The x-axis displays the tools/pipelines and the y-axis the number of TE sequences.

### TEtrimmer exhibits superior genome-wide TE annotation ability in comparison to EDTA2 and RepeatModeler2

In addition to the direct comparison of TE consensus libraries, we benchmarked EDTA2, RepeatModeler2, and TEtrimmer concerning whole-genome TE annotation by using a confusion matrix (Ou et al., 2019; Provost et al., 1998). First, all genome regions annotated as any type of TE, termed “all TEs”, were analyzed to assess the accuracy of TE masking in the genome. TEtrimmer exhibited higher sensitivity and precision (F1 scores > 0.941) than EDTA2 and RepeatModeler2 for *B. hordei*, *D. melanogaster*, *D. rerio*, and *Z. mays*, except for a slightly lower sensitivity score for the *D. rerio* EDTA2+TEtrimmer library compared to EDTA2 alone (Fig. 7A). By contrast, for *O. sativa* (rice), TEtrimmer exhibited lower sensitivity but higher precision than EDTA2 and RepeatModeler2. TEtrimmer also showed lower sensitivity and precision for *H. sapiens* compared to the other tools (Fig. 7A and Supplementary Table 5).

**Figure 7:**
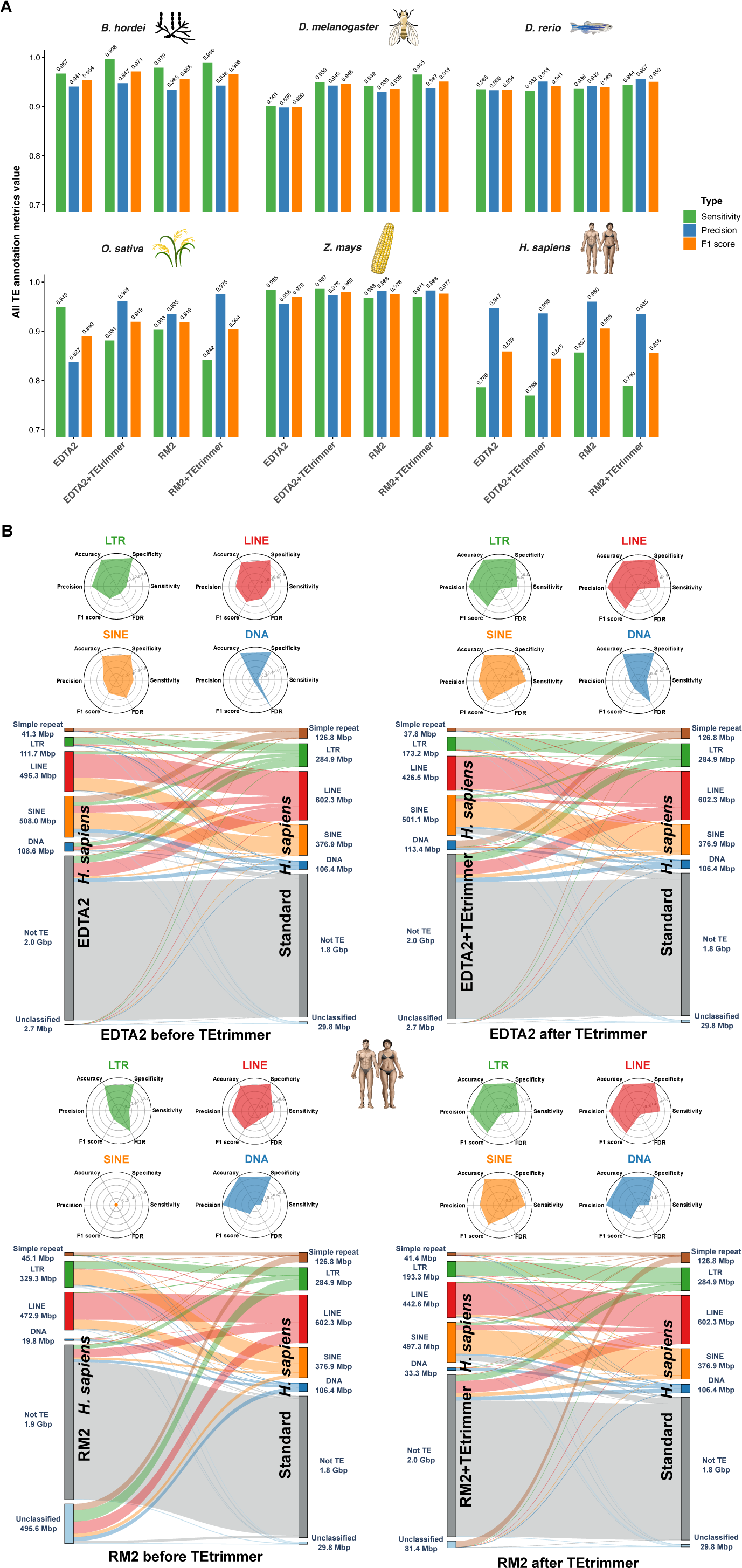
TEtrimmer exhibits superior genome-wide TE annotation ability in comparison to EDTA2 and RepeatModeler2. We assessed the genome TE annotation performance of EDTA2, RepeatModeler2 (RM2), and both after additional TEtrimmer analysis (EDTA2+TEtrimmer and RM2+TEtrimmer, respectively). All genome-wide TE annotation results based on de novo-generated libraries were compared with the results of the respective TE reference libraries (Table 2). We used a confusion matrix (Ou et al., 2019; Provost et al., 1998) to calculate the sensitivity, precision, accuracy, specificity, F1 score, and false discovery rate (FDR) of the tools. The genomes of six organisms were used for the analysis, i.e., *B. hordei* (barley powdery mildew fungus), *D. melanogaster* (fruit fly), *D. rerio* (zebrafish), *O. sativa* (rice), *Z. mays* (maize), and *H. sapiens* (human). **A** The bar plots show the sensitivity (green), precision (blue), and F1 score (orange) calculated with a confusion matrix for the six organisms indicated on top of each graph. The x-axis represents the respective analysis tools, and the y-axis displays the metrics score values based on overall genome TE annotation correctness (“All TEs”) at a range of 0.7 to 1.0. **B** Detailed genome-wide TE annotation benchmarking for *H. sapiens*. The radar plots in the upper panels show the benchmarking metrics for LTR retrotransposon (green), LINEs (red), and SINEs (orange). The lower panels display the annotation overlap and differences between de novo-generated TE libraries from the indicated tool (left) and the reference consensus TE library for *H. sapiens* (right); each bar indicates the number of Mbp masked as the respective element in the genome. The links between the left and right bars indicate the connections between the respective libraries. TE types shown are LTR retrotransposon (green), LINE (red), SINE (orange), DNA transposon (dark blue), unclassified (light blue), and genomic sequence not identified as TE (“Not TE”; grey).

Due to the overall poor performance on TE consensus library construction for *H. sapiens* by all tested tools (Fig. 6A) and TEtrimmer’s relatively lower TE annotation score for *H. sapiens* (Fig. 7A), we calculated a confusion matrix for different TE types (radar plots in Fig. 6B). In case of LTR retrotransposons, LINEs, SINEs, and DNA transposons, both the sensitivity and precision were considerably improved by additional TEtrimmer analysis. Moreover, genome-wide TE annotation based on the TEtrimmer consensus libraries exhibited better agreement with the reference annotation than those from EDTA2 and RepeatModeler2 alone (Sankey plots in Fig. 7B). For instance, the TE annotation from RepeatModeler2 (before TEtrimmer) yielded 495.6 Mbp of unclassified TEs and no SINE elements for *H. sapiens* (Fig. 7B lower panel), while additional TEtrimmer analysis reduced the unclassified TE category to 81.4 Mbp and annotated 497.3 Mbp of SINEs with high accuracy, (Fig. 7B). In conclusion, TEtrimmer libraries generally outperformed EDTA2 and RepeatModeler2-derived libraries for whole-genome TE annotation across most of the tested organisms. Additionally, TEtrimmer provided more correct TE classifications.

## Discussion

Due to decades of genome sequencing and the advanced long-read third-generation sequencing technology, the number of high-quality genomes is growing rapidly. However, precise and accurate TE annotation in genomes remains a major challenge. Several tools have been developed to automate TE discovery and annotation. Nonetheless, the manual curation of TEs is still mandatory to obtain comprehensive and detailed insights into the TE landscape of a given genome. Manual curation is a time-consuming and challenging process and often beyond the scope of a given research project. In this study, we introduce TEtrimmer, a novel software designed to automate and support the manual curation of TEs. TEtrimmer efficiently clusters and cleans MSAs and precisely defines TE boundaries. Moreover, TEtrimmer provides comprehensive report plots and a user-friendly GUI application to facilitate the inspection and improvement of discovered TE consensus sequences. In particular, the sliding window-based cleaning strategy of the MSA, which results in superior precision, and the interactive GUI application are innovative features of TEtrimmer as compared to other related software tools.

In manual curation procedures, the first labor-intensive step is separating the MSA into different clusters according to the TE sequence similarity (Fig. 2). To accelerate this step, we combined the selection of distinct MSA columns, a phylogenetic tree approach, and a machine learning method (DBSCAN) to cluster MSAs automatically and precisely. Although other clustering tools exist, none of these are fully automated. For example, TreeCluster can group phylogenetic trees according to the relative branch distance (Balaban et al., 2019), but the final cluster number must be defined manually. Furthermore, CD-HIT-EST can cluster TE sequences without generating an MSA (Fu et al., 2012), but the identity threshold needs to be manually adjusted for each MSA separately. MCHelper (Orozco-Arias et al., 2023) also provides an MSA clustering function and, like TEtrimmer, relies on DBSCAN. However, MCHelper does not include the selection of divergent MSA columns and lacks the phylogenetic tree approach. It instead uses the K2P (Kimura 2-Parameter) sequence distance matrix for DBSCAN analysis. Selectively using divergent MSA columns for clustering is crucial because consistent columns (i.e., regions of high conservation) can potentially mask or dilute the differences among sequences. Furthermore, in our experience, an un-rooted maximum likelihood phylogenetic tree is superior to calculate sequence distances for processing by DBSCAN. In contrast to the tools mentioned above, the TEtrimmer clustering algorithm automatically defines the appropriate cluster number and thresholds. It combines the selection of divergent MSA columns and a phylogenetic tree strategy for a more advanced DBSCAN clustering analysis. This approach is not only useful for the clustering of TE sequences but also for any set of nucleotide or amino acid sequences. However, because the clustering module of TEtrimmer relies on MSAs, it cannot handle large-scale datasets containing thousands of DNA sequences.

Besides MSA clustering, we developed a new MSA cleaning algorithm to remove highly divergent regions in MSAs in a specific manner (Fig. 3). Removing divergent sites from MSAs is important because they can significantly hinder the generation of TE consensus sequences and the definition of TE boundaries. Fragmented TEs and excessive end extension during the curation process typically result in the addition of many TE-unrelated flanking sequences in the MSA. This issue is especially pronounced in the case of the 5’-ends of LINE elements, which tend to be highly degraded (Goubert et al., 2022) (Fig. 3). Other tools such as trimAI (Capella-Gutiérrez et al., 2009), Clipkit (Steenwyk et al., 2020) and CIAlign (Tumescheit et al., 2022) are able to perform MSA cleaning but are mainly designed to remove gappy columns and optimize MSAs for better phylogenetic tree construction. CIAlign provides basic functions to clean rows and columns of MSAs but mainly relies on alignment gap information, which is insufficient for TE-related MSA scenarios. TEtrimmer introduces a sliding window strategy to analyze sequentially the entire MSA nucleotide by nucleotide. Our benchmarking of the cleaning performance and accuracy (Fig. 3C) demonstrates that the TEtrimmer MSA cleaning module exhibits a performance close to that of manual MSA cleaning. In summary, TEtrimmer is to the best of our knowledge the first tool that provides extensive automated MSA cleaning procedures for TE sequences.

TEtrimmer provides a powerful GUI application to facilitate the inspection and improvement of its outputs. This can close the gap between de novo-based TE libraries and the traditional, manual curation-based TE consensus libraries. TEtrimmer is the first tool to introduce a TE discovery- and curation-related GUI application. Moreover, we offer more TE-related functions in this GUI application. For example, users can not only examine and improve TEtrimmer outputs but also any TE consensus libraries established by other tools such as EDTA2 and RepeatModeler2 (Supplementary Fig. 2). They can further easily perform BLASTN searches, MSA generation, MSA sequence extension, MSA cleaning, MSA plotting, TE-Aid style plotting, and consensus sequence generation. The integration of TE benchmarking functions into the GUI application is a future option.

To test the performance of TEtrimmer regarding the discovery of intact TEs, we executed TEtrimmer using the outputs from EDTA2 and RepeatModeler2, respectively, based on six organisms of different kingdoms of eukaryotic life. The TE consensus libraries generated by TEtrimmer consistently contained a higher number of intact (“Perfect”) TEs than the consensus libraries generated by EDTA2 and RepeatModeler2 (Fig. 6A). By contrast, TEtrimmer discovered less intact LTR retrotransposons for *D. rerio*, *O. sativa* and *Z. mays* in some cases. A possible explanation for this contrasting performance could be that TEtrimmer mainly relies on the repetitive nature of TEs to improve the consensus library and, in turn, may not be effective in discovering accurately low-copy TEs. In contrast to TEtrimmer, both EDTA2 and RepeatModeler2 rely on structure-based methods to identify low-copy LTR retrotransposons. Nevertheless, such low-copy TEs can be easily recovered by TEtrimmer with the provided GUI application through the optional review and curation process.

In addition to the comparative assessment of TE integrity, we introduced a confusion matrix (Ou et al., 2019) to evaluate the TE annotation performance at a genome-wide scale. EDTA2 and RepeatModeler2 exhibited great sensitivity and precision for the selected organisms at the level of “all TEs”, which indicates that these tools can accurately mask TE regions in the respective genomes. However, most of the benchmarking metrics were improved after TEtrimmer analysis, except for *O. sativa* and *H. sapiens* (Fig. 7A) where the metrics values were slightly decreased, most likely due to the relatively poor performance of TEtrimmer regarding the analysis of low-copy TEs. Nevertheless, we found an overall improved TE classification accuracy for *H. sapiens* by TEtrimmer (Fig. 7B). Here, we introduced for the first time Sankey plots to the TE field to visualize effectively the quality of TE annotation compared to the respective reference library.

To date, EarlGrey and MCHelper are the two most prominent tools designed for automating the manual curation of TEs. TEtrimmer outperforms these tools in five main aspects. (1) EarlGrey does not provide a clustering function for MSAs, which can be a limitation for analyzing highly divergent TEs. Compared to MCHelper, TEtrimmer has a more sophisticated MSA clustering strategy that is based on the selection of divergent MSA columns in combination with the generation of a phylogenetic tree, which can dramatically improve DBSCAN performance. (2) Both EarlGrey and MCHelper are not able to clean efficiently MSAs, due to a lack of functions to clean MSA rows. (3) For each iteration of MSA extension, EarlGrey and MCHelper extend all TE sequences in the MSA. By contrast, we developed an MSA extension algorithm that can selectively extend TE sequences based on the divergence in the alignment. This strategy facilitates even the analysis of highly degraded regions, such as the 5’ regions of LINE elements. (4) TEtrimmer assigns different evaluation levels for each output based on criteria such as the presence of terminal repeats, classification status, MSA sequence number, number of full-length BLASTN hits, and PFAM domain prediction (Table 1). This helps in systematically categorizing and assessing the quality of the outputs. (5) TEtrimmer provides comprehensive report plots (Fig. 4) and a user-friendly GUI application (Fig. 5), which allow users to inspect and improve each TEtrimmer output sequence conveniently. We also integrated the MSA cleaning, MSA extension, and plotting functions into the GUI application to achieve manual curation-level TE libraries efficiently.

### Conclusions

In conclusion, TEtrimmer is a powerful and user-friendly tool designed to automate the manual curation of TEs, addressing the major challenge of accurately annotating these repetitive DNA sequences in eukaryotic genomes. By integrating advanced techniques for MSA clustering, cleaning, and extension, as well as TE boundary definition, TEtrimmer enhances the quality of TE annotations markedly. The tool provides detailed report plots and a GUI application for efficient inspection and refinement of results, making it accessible even for researchers that lack extensive levels of expertise in TE genetics. Comprehensive benchmarking against the genomes of six diverse eukaryotic organisms demonstrated the advanced performance of TEtrimmer compared to existing tools like RepeatModeler2 and EDTA2, particularly in identifying intact (full-length) TEs. Hence, TEtrimmer bridges the gap between automated TE annotation and the gold standard of manual curation, offering a robust solution for the accurate and efficient annotation of TEs in genomic studies.

## Methods

### BLASTN searches

TEtrimmer (v1.3.0) uses the TE consensus library output from any de novo TE discovery tool such as EDTA2, RepeatModeler2, and REPET, or consensus TE libraries from closely related species, as input to improve the TE library quality. First, TEtrimmer removes duplicated input sequences via CD-HIT-EST (Fu et al., 2012) (v.4.8.1) when the “`--dedup`” option is enabled. Sequences with more than 95% sequence identity and coverage are merged into one element. Then, each input sequence is converted into a single file to enable multi-thread analysis. TEtrimmer uses these separated sequences to perform BLASTN searches against the respective genome to find query copies for each TE sequence. By default, only the 70 longest and 30 randomly selected BLASTN hits are used for further analysis when more than 100 target sequences are identified by the BLASTN query, as these are typically sufficient for meaningful downstream analysis. Finally, the BLASTN search result is converted into a BED file.

### MSA generation

TE sequences are extracted based on the BED file derived from the BLASTN search of each input TE consensus sequence against the corresponding genome (see above) by “`bedtools getfasta`” (Quinlan & Hall, 2010) (v2.31.1). Those sequences are aligned by MAFFT (Katoh & Standley, 2013) (v7.520) with the fast alignment option “`--retree 1`” and any gappy alignment columns are removed prior to MSA clustering by the function “`remove_gap_columns`” (see section “MSA cleaning” below).

### MSA clustering

Because of the typically low sequence divergence among TE subfamilies, several types of sub-families are usually included in the MSA based on the BLASTN search derived from each input consensus sequence against the respective genome (Fig. 2). To separate these subfamilies, TEtrimmer calculates the percentage of each type of nucleotide in every column. Gaps in that column are not included in the calculation. Divergent columns are defined as those where all nucleotide proportions are lower 0.8 in that column. All divergent columns are merged together into a new FASTA file, yielding an MSA of pseudo-sequences representing the divergent regions in the original MSA, which is used by IQ-TREE (Minh et al., 2020) to generate a maximum likelihood phylogenetic tree. We use the fixed IQ-TREE model “K2P+I” to reduce the execution time of this module. Then, a sequence distance matrix is calculated based on the relative tree branch distances. Finally, the DBSCAN (Birant & Kut, 2007) machine learning method is performed based on the sequence distance matrix to cluster the sequences of the MSA, with epsilon set to 0.1 and minimum samples set to 2. More than two clusters can be generated after MSA clustering. By default, TEtrimmer only proceeds with the top two clusters containing the highest number of sequences, which provides in our hands typically a good balance between sensitivity and computational intensity. Users can increase this number by the option “`--max_cluster_num`” but need to consider that increasing the cluster number can cause a high rate of false-positive results (i.e., non-TE sequences mistakenly classified as TEs).

### MSA cleaning (“trimming”)

For cleaning of columns within an MSA, TEtrimmer uses the function “`remove_gap_columns`”. By default, TEtrimmer removes columns when the gap comprises more than 80% of the column or the nucleotide number in this column is less than five. In a subsequent step, columns with a gap percentage between 40% and 80% and where the predominant nucleotide constitutes less than 70% of the total nucleotides are deleted.

In addition to gappy columns, highly divergent sequence regions might occur in the MSA. TEtrimmer includes two functions to remove divergent regions from the ends of the MSA row by row. The function “`crop_end_by_divergence`” first calculates the proportion of each nucleotide in every column that have at least five nucleotides. Gaps are not included in this calculation. All nucleotide proportions in columns with fewer than five nucleotides are considered as zero. Then the entire MSA is converted into a proportion matrix by replacing each nucleotide with the corresponding proportion value and all gap positions are converted to zero. For each row of the proportion matrix, two sliding windows (default size 40 bp) are created at both ends that move stepwise towards the opposite end of the sequence nucleotide position by nucleotide position. Each sliding window stops when the mean of the nucleotide proportions within the sliding window is greater than the set threshold (default 0.7), indicative of having reached a conserved region. For each sliding window starting from the left end of the sequence, all nucleotide positions between the left end of the sequence and the stopping point (left boundary) of this sliding window are deleted. Similarly, for each sliding window that starts from the right end of the sequence, all nucleotide positions between the right end of the sequence and the stopping point (right boundary) of the sliding window are deleted. After completion of all rows, the process is repeated with a smaller sliding window (default size 4 bp) and a higher threshold (default 1.0) for fine-polishing the MSA ends.

The second function used to clean the MSA is called “`crop_end_by_gap`”. It deploys a similar sliding window cleaning principle as “`crop_end_by_divergence`”, but the difference is that “`crop_end_by_gap`” will not convert the MSA into a proportion matrix. Instead, it takes advantage of the original MSA. For each sequence in the MSA, two sliding windows (default size 250 bp) are created at both ends that stepwise move towards the opposite end of the sequence nucleotide position by nucleotide position. Each sliding window stops when the proportion of gaps included in the sliding window is below the set threshold (default 0.1), indicative of having reached a conserved region. For each sliding window that starts from the left end of the sequence, all nucleotide positions between the left end of the sequence and the stopping point (left boundary) of this sliding window are deleted. Similarly, for each sliding window that starts from the right end of the sequence, all nucleotide positions between the right end of the sequence and the stopping point (right boundary) of the sliding windows are deleted. The “`crop_end_by_gap`” function is also used to facilitate MSA extensions (see below).

### MSA sequence extension

Annotated TEs derived from de novo TE discovery software can be fragmented and truncated. To complete such TE sequences, TEtrimmer iteratively extends both ends of the sequences from the clustered MSA separately via “`bedtools slop`” (Quinlan & Hall, 2010). The default step size is 1000 bp for each end per extension, with a maximum total extension size of 7000 bp at each end of the sequences.

First, TEtrimmer performs left end sequence extension of the MSA. To save computing time, after each round of sequence extension TEtrimmer only aligns the newly extended sequences from the current round, followed by cleaning of gappy columns via the function “`remove_gap_columns`” as described above (see section “MSA cleaning”). Then, the function “`crop_end_by_gap`” (see section “MSA cleaning” above) is employed to clean the MSA additionally row by row, and the sum of the removed sites for each sequence (row) is recorded. The sequence is excluded from the next round of extension when the sum of its removed sites by “`crop_end_by_gap`” is more than 90% of the sequence length, indicating a sufficient extension for this sequence as seemingly flanking genomic regions have been reached. Moreover, to assess if the overall left end extension is sufficient to include the left TE boundary, the newly extended and cleaned MSA (only containing the sequences extended in the current round) is used to generate a consensus sequence where the letter “N” represents any ambiguous position. An ambiguous position is an MSA column site where the proportion of the predominant nucleotide is below 0.7 or the total nucleotide count is below five. Then, a sliding window (default size 150 bp) is created at the left end of the TE consensus sequence that moves stepwise towards the opposite end of the sequence nucleotide position by nucleotide position. The sliding window stops when the proportion of the ambiguous letter “N” (as defined above) within the sliding window is less than the set threshold (default 0.3). If the stopping point (left boundary) of the sliding windows is more than 300 bp away from the left end of the consensus sequence, TEtrimmer regards the left end sequence extension as complete. Otherwise, a new round of extension is initiated.

When the left end sequence extension is finished, TEtrimmer conducts right end sequence extension of the MSA. Similar to the steps described above, right end sequence extension also only aligns extended sequences from the current round and generates the respective consensus sequence. In contrast to the left end, to judge if the overall right end extension is sufficient, a right end sliding window is created only for this consensus sequence. The sliding window stops when the proportion of the ambiguous letter “N” (as defined above) within the sliding window is less than the set threshold (default 0.3). If the stopping point (right boundary) of the sliding window is more than 300 bp away from the right end of the consensus sequence, the right end extension is complete.

Note that this way extension sizes for the two ends of each sequence in the MSA may differ. After sequence extension at both ends, TEtrimmer calculates the new (extended) genome coordinates (locations) for each sequence. Finaly, extended sequences are extracted from the corresponding genome. All sequences are aligned and gappy columns are removed from the MSA by the function “`remove_gap_columns`” as described above (see section “MSA cleaning”).

### TE boundary definition

Sequence extension helps to complete TE consensus sequences, but putative excessive extension can also add other, non-TE sequences (false-positives) at both ends of the MSA. TEtrimmer uses two strategies to identify TE boundaries and to remove such inappropriate regions. Based on the final MSA after the extension step, first a consensus sequence is generated for each MSA, and “N” is used to represent any ambiguous column where the proportion of the predominant nucleotide is below 0.8 or the total count of nucleotides is less than five. Then, a self-BLASTN search (with relatively relaxed e-value of 0.05) of the consensus sequence is conducted to identify potential terminal repeats of TEs, including LTRs and TIRs. Once a terminal repeat is found, the TE boundary is set at the distal position of the terminal repeat, and the MSA is cropped accordingly. On the other hand, if no terminal repeat is identified, TEtrimmer selects sliding windows (default size 150 bp) at both ends of the consensus sequence that stepwise move towards the opposite end of the sequence nucleotide position by nucleotide position until the proportion of the ambiguous letter “N” is below 0.2 within the sliding windows. Then, only the MSA columns between the stopping points of the two sliding windows (including the sliding windows) are kept. This procedure removes the majority of false-positive flanking sequences. Furthermore, the functions “`crop_end_by_divergence`” and “`crop_end_by_gap`” (see section “MSA cleaning above) remove highly divergent regions from the MSA. Based on the defined TE boundaries, TEtrimmer generates the corresponding consensus sequence for further analysis.

### ORF and PFAM domain prediction

For autonomous TEs, the presence of characteristic protein domains can be used to assist TE classification (Goubert et al., 2022) and to determine TE orientation. TEtrimmer adopts the PFAM database (Mistry et al., 2021) to annotate protein domains within TEs. It either uses the “`--pfam_dir`” option to query a local PFAM library, or it can download the current PFAM database automatically in the absence of a local PFAM instance. Thereafter, TEtrimmer uses the EMBOSS function “`getorf`” (Rice et al., 2000) to predict ORFs and “`pfam_scan.pl`” (Mistry et al., 2021) to search for TE protein domains against the PFAM database. Typically, all predicted PFAM domains share the same orientation. TEtrimmer generates reverse complements of the TE consensus sequence and the MSA if all PFAM domains are found on the antisense strand of the consensus sequence. If TE-related PFAM domains are found on both strands, TEtrimmer sums the length of PFAM domains found on both strands to decide on the orientation of the TE consensus sequence via a majority decision.

### TE classification

TEtrimmer includes two TE classification methods. First, it adopts the RepeatModeler2 module “`RepeatClassifier`” (Flynn et al., 2020) to categorize TEs. By default, TEtrimmer re-assigns processed TEs if the sequence extension size is larger than 4000 bp as these may represent mis-categorized and/or fragmented TEs. Further, two options are associated with the first classification method, named “`--classify_all`” and “`--classify_unknown`”. TEtrimmer runs RepeatClassifier for each output if the “`--classify_all`” option is used and only re-classifies “Unknown” TEs via the “`--classify_unknown`” option. If RepeatClassifier assigns a new category, TEtrimmer will replace the old assignment with it; alternatively, the original classification is kept. The second method is performed when the initial analysis of all the input TE sequences is completed. TEtrimmer uses the final, classified output sequences as a database to re-assign unknown sequences via RepeatMasker.

### Output evaluation

Five different evaluation levels including “Perfect”, “Good”, “Reco_check” (i.e., recommended check), “Need_check”, and “Low_copy” are assigned to each TEtrimmer output. TEtrimmer evaluates TE consensus sequences based on the presence of terminal repeats, classification status, the number of sequences in the MSA, the number of full-length BLASTN hits, and PFAM protein domain predictions (Table 1). A full-length BLASTN hit is defined as a sequence with more than 90% coverage and more than 85% identity with the respective query sequence (i.e., the TEtrimmer-processed consensus sequence). When the number of sequences within the MSA is less than ten, TEtrimmer examines if the input sequence contained terminal repeats and if the number of full-length BLASTN hits is greater than or equal to two. If so, this input sequence is regarded as a low-copy element, otherwise it is excluded from further analysis.

### TE consensus library de-duplication

All consensus sequences processed by TEtrimmer are stored into the file “TEtrimmer_consensus.fasta”, which potentially contains duplicated sequences. To reduce any putative redundance, TEtrimmer performs two rounds of de-duplication by CD-HIT-EST (Fu et al., 2012). For the first round, consensus sequences are grouped into one cluster if they share more than 90% sequence identity and the alignment coverage of the shorter sequence is more than 90%. Afterwards, TEtrimmer selects all clusters that contain at least one “Perfect” or “Good” (Table 1) consensus sequence. For each selected cluster, if “Perfect” candidates exist, the longest “Perfect” consensus sequence is selected as the representative. If only “Good” candidates are present, the longest “Good” consensus sequence is selected. The second de-duplication round extracts all sequences from the remaining clusters and uses these sequences again as input for CD-HIT-EST. Sequences are grouped into one cluster if they have more than 85% identity and the alignment coverage for all sequences is greater than 80%. Only the longest sequence in each new cluster is selected for the de-duplicated TE consensus library. The de-duplicated TE consensus sequences are saved into the text file “TEtrimmer_consensus_merged.fasta”.

### Generation of report plots

Seven plots are generated for each output to help evaluating the TEtrimmer output and to compare the differences between the TEtrimmer-processed and input consensus sequences. The report plots file include (1) an MSA ends plot, (2) a whole MSA plot, (3 and 4) TE-Aid (Goubert et al., 2022) plots for both input and output sequences, (5 and 6) a PFAM plot for both input and output sequences, and (7) a dot plot comparing input and output sequences (Fig. 7). For the MSA ends plot, TEtrimmer extracts 50 columns before and after the identified TE boundaries (see section “TE boundary definition” above) from the MSA and artifactually joins them by ten hyphen symbols, signifying the TE (Fig. 4A). Nucleotides are represented by different colors, and the nucleotide background is colored if the nucleotide proportion in the column is less than 40%. The whole MSA plot module is derived from the software CIAlign (Tumescheit et al., 2022) and the nucleotides are represented by different colored bars (Fig. 4B). In contrast to the default CIAlign plot, TEtrimmer adds two arrows to indicate the TE boundaries (TE start and TE end) in the whole MSA plot. TEtrimmer also includes the TE-Aid package to help plotting. All TEtrimmer output and input TE consensus sequences are used for TE-Aid analysis (Fig. 4C and D). Furthermore, the PFAM plots are generated based on ORF and PFAM domain prediction results, which are denoted as blue and red arrows, respectively (Fig. 4E). The last dot plot is generated by the EMBOSS tool “`dotmatcher`” (Rice et al., 2000). The position of the consensus output sequence after analysis by TEtrimmer is plotted on the x-axis and the position of the corresponding input TE consensus sequence is plotted on the y-axis of the dot plot (Fig. 4F). All seven report plots are merged into one file to be used for convenient output evaluation.

### Used genomes and reference TE consensus libraries for genome-wide benchmarking

The genomes of six eukaryotic organisms were used to test the TE annotation performance of TEtrimmer at the level of TE consensus libraries and a genome-wide scale. These species comprised *B. hordei* (barley powdery mildew fungus), *D. melanogaster* (fruit fly), *D. rerio* (zebrafish), *O. sativa* (rice), *Z. mays* (maize), and *H. sapiens* (human) (Table 1). The reference TE consensus library of *B. hordei* is based on in-house manually curated TEs (Qian et al., 2023). In case of *D. melanogaster*, *D. rerio*, *O. sativa*, and *Z. mays*, Repbase 28.10 (Weidong. Bao et al., 2015) was used for generating the respective reference TE consensus libraries. *H. sapiens* reference TE consensus library was extracted from Dfam (Hubley et al., 2016).

### Benchmarking of MSA cleaning performance

Due to the presence of fragmented TEs and/or excessive end extension, the extended MSAs typically contain regions of low sequence conservation, which can be efficiently cleaned by the TEtrimmer cleaning module. The MSA cleaning capability of TEtrimmer was evaluated quantitatively by a customized confusion matrix (Provost et al., 1998). Three MSAs were used for each evaluation, including the original MSA before cleaning, the manually cleaned MSA, and the MSA cleaned by TEtrimmer. The manually cleaned MSA served as the reference in the benchmarking process.

Both manual cleaning and TEtrimmer-based cleaning did not delete the nucleotides in lowly conserved (poorly aligned) regions but converted them to gaps (-). Consequently, all three MSAs had the same number of columns and sequences. The order of the sequences within the MSA was also identical. Thereafter, each position in the three MSAs was analyzed and corresponding true positive (TP) sites, false negative (FN) sites, false positive (FP) sites, and true negative (TN) sites values were calculated and summed into TP, FN, FP, and TN values (Table 3). For example, if a site in the original MSA was occupied by any type of nucleotide, this site was counted as “true positive” when both manual and TEtrimmer-based cleaning converted this site to a gap (-). At the end, the TEtrimmer cleaning sensitivity (TP / (TP + FN)), precision (TP / (TP + FP)), and F1 score (2 * (sensitivity * precision) / (sensitivity + precision)) were calculated and used to evaluate the TEtrimmer cleaning performance.

**Table 3.**
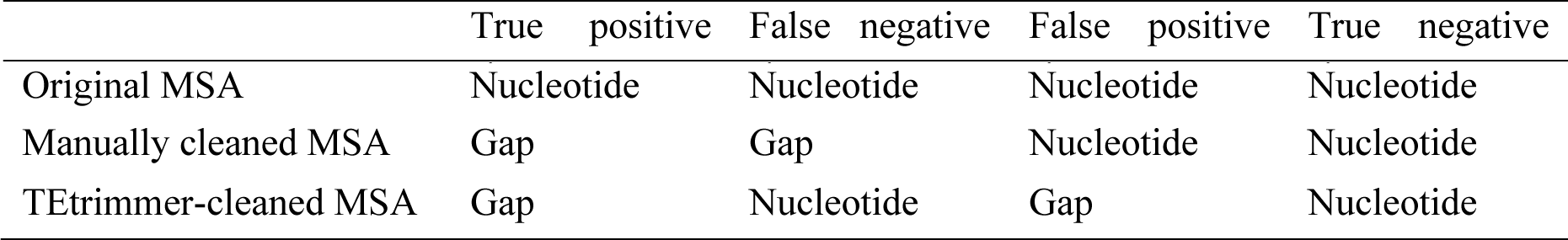
Confusion matrix calculation for the MSA cleaning performance.

|  | True positive | False negative | False positive | True negative |
| --- | --- | --- | --- | --- |
| Original MSA | Nucleotide | Nucleotide | Nucleotide | Nucleotide |
| Manually cleaned MSA | Gap | Gap | Nucleotide | Nucleotide |
| TEtrimmer-cleaned MSA | Gap | Nucleotide | Gap | Nucleotide |

### Benchmarking of genome-wide TE discovery and annotation

We benchmarked TEtrimmer, RepeatModeler2, and EDTA2 on the basis of the corresponding reference TE libraries (Table 1) by two methods. The first method directly compared the consensus sequences to assess the integrity of TE sequences, which aimed to determine how many TE sequences from the reference library can be recovered by the respective de novo-generated library. The other method compared genome-wide TE annotation by the different tools to evaluate if TEs in the genome were correctly masked. Six organisms were used for the benchmarking test and the reference TE consensus libraries and genomes were downloaded and prepared (Table 1). RepeatModeler2 (Flynn et al., 2020) with parameters `“-threads 50 -LTRStruct`” and EDTA2 (https://github.com/oushujun/EDTA/tree/EDTA2) with parameters “`--threads 50 -- sensitive 1`” were run to generate de novo TE libraries for all selected organisms. Thereafter, de novo-generated TE consensus libraries were used as the input for TEtrimmer analysis with parameters “`--num_threads 50 --classify_all`”. Finally, all TE consensus libraries generated before and after TEtrimmer analysis were used individually to annotate the corresponding genome by RepeatMasker v4.1.1 with parameters “`-pa 50 -s -a -inv -gff`”.

First, we performed benchmarking by direct comparison of the TE consensus libraries. We used the benchmarking method developed by Flynn and co-workers (Flynn et al., 2020) for this purpose. All TE consensus libraries generated by EDTA2, RepeatModeler2, and TEtrimmer were compared with the corresponding reference TE consensus library (Table 1). For EDTA2, we created an additional library for the benchmarking by combing the EDTA2 consensus library with all TE sequences identified by the EDTA2 structure-based module. This new EDTA2 library was then de-duplicated using CD-EST-EST with the parameters “`-c 0.9 -aL 0.9 -aS 0.9`”, resulting in the new library named “EDTA2 struct-lib”. Due to the default separation of flanking LTRs from internal sequences (LTR-INT) in the case of LTR retrotransposon by tools like EDTA2 and RepeatModeler2, as well as in reference TE consensus libraries, a custom Python script was developed to separate flanking LTRs from the internal sequences (LTR-INT) for all TEtrimmer consensus libraries. The respective script was named “SeparateLTR.py”, which can be downloaded from the TEtrimmer GitHub repository. To avoid the benchmarking bias caused by flanking LTRs, we did not include the flanking LTRs for benchmarking. Four evaluation levels were assigned to each TE sequence in the reference library, including “Perfect”, “Good”, “Present”, and “Not found”. “Perfect” means that the reference TE consensus sequence has one match in the de novo-generated library with > 95% similarity and coverage. “Good” indicates that the reference TE consensus sequence has multiple overlapping matches in the de novo-generated library, each with > 95% similarity and total coverage. “Present” is similar to “Good”, but the required minimum similarity and total coverage is decreased to 80%.

Next, we performed the benchmarking at the level of genome-wide TE annotation. We developed a custom python script named “TE_sankey_plotter.py” (available from TEtrimmer GitHub repository), which employs a confusion matrix to evaluate the genome-wide TE annotation and to illustrate the outcome comparatively via Sankey plots. The details about how to apply a confusion matrix on the genome-wide TE annotation was described before (Ou et al., 2019). Briefly, the RepeatMasker “.out” annotation files derived from the TEtrimmer, RepeatModeler2, and EDTA2 consensus libraries, respectively, were compared with the RepeatMasker “.out” file based on the reference TE consensus library. First, all “.out” files were sequentially converted into BED files with four columns (chromosome name, start position, end position, and TE type). Then, the overlapping TE regions were resolved by BEDtools v2.31.1 (Quinlan & Hall, 2010) with parameters “`-d 0 -c 4,4,2,3 -o collapse, distinct, collapse, collapse`”. Finally, six metrics scores were calculated, i.e., sensitivity, specificity, accuracy, precision, F1 score, and false discovery rate, to evaluate the genome-wide TE annotation.

### Computational resource requirements and installation of TEtrimmer

TEtrimmer is written in the language Python3 and can be conveniently installed via a Conda package or a Docker image at https://github.com/qjiangzhao/TEtrimmer. TEtrimmer can be run on the operational systems Linux, macOS, and a Windows subsystem for Linux (WSL). There are no specific hardware requirements for running TEtrimmer. TEtrimmer is multi-threaded tool and supports high-performance computing (HPC).

## Abbreviations

bp: base pair
DBSCAN: Density-Based Spatial Clustering of Applications with Noise
EDTA2: Extensive De novo Transposon Annotation 2
GUI: Graphical user interface
LINE: Long interspersed nuclear element
LTR: Long terminal repeat
LTR-INT: Long terminal repeat internal sequence
MSA: Multiple sequence alignment
ORF: Open reading frame
PCA: Principal component analysis
PFAM: Protein families database
SINE: Short interspersed nuclear element
TE: Transposable element
TIR: Terminal inverted repeat

## Declarations

### Acknowledgments

We would like to thank to Anna V. Protasio, Tony Heitkam, and Lera Emmanuelle for their valuable feedback on the TEtrimmer software. We acknowledge Xinyi Liu (TUM, Munich, Germany) pre-testing TEtrimmer. We are grateful to Clément Goubert (University of Arizona, USA), who allowed the addition of TE-Aid into the TEtrimmer package. The TEtrimmer development and benchmarking was supported by high performance computing at RWTH Aachen University under project IDs rwth0146 and rwth1554.

### Author contributions

RP, SK, and JQ conceived this study.

JQ and HX developed the TEtrimmer software.

SK performed proofreading of the TEtrimmer code.

HX and JQ drafted the TEtrimmer manual.

LF supported the development of the TEtrimmer MSA cleaning procedure and the MSA clustering algorithm.

SO provided EDTA2 results and performed intensive de-bugging of TEtrimmer.

JS conducted RepeatMasker analysis.

JQ and SK wrote the first draft of the manuscript.

JQ drafted the Figures.

MW supervised work performed by HX and provided funding.

RP and SK supervised work performed by JQ.

RP provided funding.

All authors reviewed the manuscript and contributed to the editing.

### Ethics approval

Not applicable.

### Data availability

TEtrimmer is an open-source software under license GPLv3. We used large language models ChatGPT and GPT4 to facilitate the development of TEtrimmer. The code can be downloaded from GitHub at https://github.com/qjiangzhao/TEtrimmer. A detailed TEtrimmer manual is supplied. All benchmarking results and scripts can be found at GitHub at https://github.com/qjiangzhao/TEtrimmerPaperFile. All third-party software included in the TEtrimmer package are published under open-source licenses, with the exception of TE-Aid, for which we have approval by the author Clément Goubert (University of Arizona, USA).

## Supplementary files

**Supplementary Figure 1.** Multiple sequence dot plot based on all sequences in a TE cluster.

**Supplementary Figure 2.** The TEtrimmer GUI application can help to inspect and improve any TE consensus library.

**Supplementary Table 1.** Relative tree branch distance matrix of the un-rooted maximum-likelihood phylogenetic tree calculated on the basis of the divergent columns in the MSA.

**Supplementary Table 2.** An exemplary report table generated by TEtrimmer.

**Supplementary Table 3.** Number of TE sequences in each benchmarking category in comparison to the respective reference TE consensus library (raw data for Fig. 6A).

**Supplementary Table 4.** Types of TEs in the “Perfect” benchmarking class (raw data for Fig. 6B).

**Supplementary Table 5.** Confusion matrix scores related to the genome-wide benchmarking.

**Supplementary File 1.** Example of interactive report plots generated by the GUI application.

